# The mesoglea buffers the physico-chemical microenvironment of photosymbionts in the upside-down jellyfish *Cassiopea* sp

**DOI:** 10.1101/2022.10.06.511124

**Authors:** Niclas Heidelberg Lyndby, Margaret Caitlyn Murray, Erik Trampe, Anders Meibom, Michael Kühl

## Abstract

The jellyfish *Cassiopea* has a conspicuous lifestyle, positioning itself upside-down on sediments in shallow waters thereby exposing its photosynthetic endosymbionts (Symbiodiniaceae) to light. Several studies have shown how the photosymbionts benefit the jellyfish host in terms of nutrition and O_2_ availability, but little is known about the internal physico-chemical microenvironment of *Cassiopea* during light-dark periods. Here, we used fiber-optic sensors to investigate how light is modulated at the water-tissue interface of *Cassiopea* sp. and how light is scattered inside host tissue. We additionally used electrochemical and fiber-optic microsensors to investigate the dynamics of O_2_ and pH in response to changes in the light availability in intact living specimens of *Cassiopea* sp.

Mapping of photon scalar irradiance revealed a distinct spatial heterogeneity over different anatomical structures of the host, where oral arms and the manubrium had overall higher light availability, while shaded parts underneath the oral arms and the bell had less light available. White host pigmentation, especially in the bell tissue, showed higher light availability relative to similar bell tissue without white pigmentation. Microprofiles of scalar irradiance into white pigmented bell tissue showed intense light scattering and enhanced light penetration, while light was rapidly attenuated over the upper 0.5 mm in tissue with symbionts only.

Depth profiles of O_2_ concentration into bell tissue of intact, healthy/living jellyfish showed increasing concentration with depth into the mesoglea, with no apparent saturation point during light periods. O_2_ was slowly depleted in the mesoglea in darkness, and O_2_ concentration remained higher than ambient water in large (> 6 cm diameter) individuals, even after 50 min in darkness. Light-dark shifts in large medusae showed that the mesoglea slowly turns from a net sink during photoperiods into a net source of O_2_ during darkness. In contrast, small medusae showed a more dramatic change in O_2_ concentration, with rapid O_2_ buildup/consumption in response to light-dark shifts; in a manner similar to corals. These effects on O_2_ production/consumption were also reflected in moderate pH fluctuations within the mesoglea. The mesoglea thus buffers O_2_ and pH dynamics during dark-periods.

## Introduction

Symbiont-bearing jellyfish in the genus *Cassiopea* exhibit a conspicuous lifestyle by settling upside-down on sediment, exposing the subumbrella and oral arms to light. The oral side (i.e., the subumbrella side) of the medusa is particularly dense in intracellular microalgal symbionts, which are dinoflagellates of the family Symbiodiniaceae (Lampert, 2016). In contrast to corals where symbionts are found in endoderm cells, symbiont algae in *Cassiopea* are kept in clusters in host-specialized amoebocyte cells, directly below the epidermis in the mesoglea of the bell and oral arms (Colley and Trench, 1985; Estes et al., 2003). Similar to many photosymbiotic corals and anemones, *Cassiopea* rely on a metabolic exchange of carbon and nitrogen between the host animal and its algal symbionts (Welsh et al., 2009; Freeman et al., 2016; Lyndby et al., 2020). Translocation of autotrophically acquired carbon from the algal symbionts to the host covers a significant part of its daily carbon requirement, estimated to be at a similar or greater level than what is known from reef-building corals (Muscatine et al., 1981; Verde and McCloskey, 1998).

While most photosymbiotic cnidarians require a rather stable environment to maintain a healthy symbiosis, *Cassiopea* appears extraordinarily resilient to fluctuating environmental conditions (Goldfarb, 1914; Morandini et al., 2017; Aljbour et al., 2019; Klein et al., 2019; Banha et al., 2020) and are increasingly regarded as an invasive species (Mills, 2001; Morandini et al., 2017). *Cassiopea* are typically found in shallow waters (e.g. lagoons, around seagrass beds, and mangroves) in tropical and subtropical regions (Drew, 1972; Hofmann et al., 1996) that are prone to strong diel fluctuations in both temperature, and salinity, as well as high solar irradiance (Anthony and Hoegh-Guldberg, 2003; Veal et al., 2010). Human activities in such coastal ecosystems can further add to strong fluctuations in both nutrient input, O_2_ availability, and pH (Stoner et al., 2011; Klein et al., 2017; Rowen et al., 2017; Arossa et al., 2021).

The photobiology and optical properties of marine symbiont-bearing cnidarians have been studied in detail in reef-building corals, showing adaptations in host growth patterns, microscale holobiont light modulation, and colony-wide symbiont organization to enable and optimize symbiont photosynthesis for the benefit of the host. There has been a special focus on coral skeletons and host pigments (e.g. Falkowski et al., 1984; Kühl et al., 1995; Wangpraseurt et al., 2014b; Lyndby et al., 2019; Kramer et al., 2021; Bollati et al., 2022). Studies have shown that *in vivo* light exposure of symbionts is modulated on a holobiont level to enhance photosynthesis, including alleviating photodamage and bleaching (Lesser and Farrell, 2004; Enriquez et al., 2005; Marcelino et al., 2013; Wangpraseurt et al., 2017), altering inter- and intracellular pH (e.g. Kühl et al., 1995; Gibbin et al., 2014), and directly affecting holobiont temperature (Jimenez et al., 2008, 2012; Lyndby et al., 2019). Recently, these experimental insights were integrated in a first multiphysics model of radiative, heat and mass transfer in corals, simulating how coral tissue structure and composition can modulate the internal light, O_2_ and temperature microenvironment of corals (Taylor Parkins et al., 2021). Given a similar global distribution and a benthic lifestyle relying on host-symbiont metabolic interactions, it is feasible to assume that *Cassiopea* exhibit similar traits, but few studies have investigated to what extend *Cassiopea* modulate their physicochemical microenvironment (Klein et al., 2017; Arossa et al., 2021).

In this study, we use a combination of optical and electro-chemical microsensors to investigate the internal microenvironment of *Cassiopea* sp. medusae. We explore how light is modulated by host morphology and distinct anatomical features. Furthermore, we show how the size of animals and the thickness of their bell mesoglea affects the intracellular physicochemical microenvironment of intact, living *Cassiopea* sp. medusae.

## Methods

### *Cassiopea* maintenance

Medusae of *Cassiopea* sp. were acquired via DeJong Marinelife (Netherlands) in 2018, and since cultivated at the aquarium facility at the Marine Biology Section in Helsingør, University of Copenhagen (Denmark). According to the commercial provider, the obtained specimens originated from Cuba indicating the species *Cassiopea xamachana* or *C. frondosa*, but no species identification was performed. The medusae were kept in artificial seawater (ASW; 25°C, 35 ppt, pH of 8.1) in a 60 L glass aquarium, under a 12h-12h day-night cycle using LED lamps (Tetra, Pacific Sun) providing an incident photon irradiance (400-700 nm) of ~300 µmol photons m^−2^ s^−1^, as measured with a cosine corrected mini quantum sensor (MQS-B, Walz, Germany), connected to a calibrated irradiance meter (ULM-500, Walz, Germany). Animals were fed with living *Artemia* sp. nauplii twice a week, and 25% of the aquarium water was replaced with fresh filtered ASW (FASW) 1 hour after feeding. Additionally, water was continuously filtered using an internal filter pump, a filter sock (200 µm), and a UV filter. Additionally, an air pump was used to ensure water remained oxygenated at all times.

### Light measurements

#### Photon scalar irradiance

Mapping of photon scalar irradiance in µmol photons m^−2^ s^−1^ (PAR; 400–700 nm) on the jellyfish tissue surface was performed with a submersible spherical micro quantum sensor (3.7 mm diameter; US-SQS/L, Walz, Germany) connected to a calibrated irradiance meter (ULM-500, Walz, Germany). Measurements were conducted inside an aquarium tank filled with ASW and the inside covered with a black cloth to avoid internal reflections from the container. The medusa was placed in the aquarium and illuminated vertically from above by a fiber-optic tungsten-halogen lamp (KL-2500 LCD, Schott GmbH, Germany), equipped with a collimating lens. Photon scalar irradiance was measured holding the sensor at a ~45° angle, while measuring light on the surface of the animal within 7 areas of interest (AOI) of one animal (Figure 1B). Measurements were normalized against the incident downwelling photon irradiance, as measured over the black cloth covering the inside of the container relative to each AOI on the animal.

**Figure 1.**
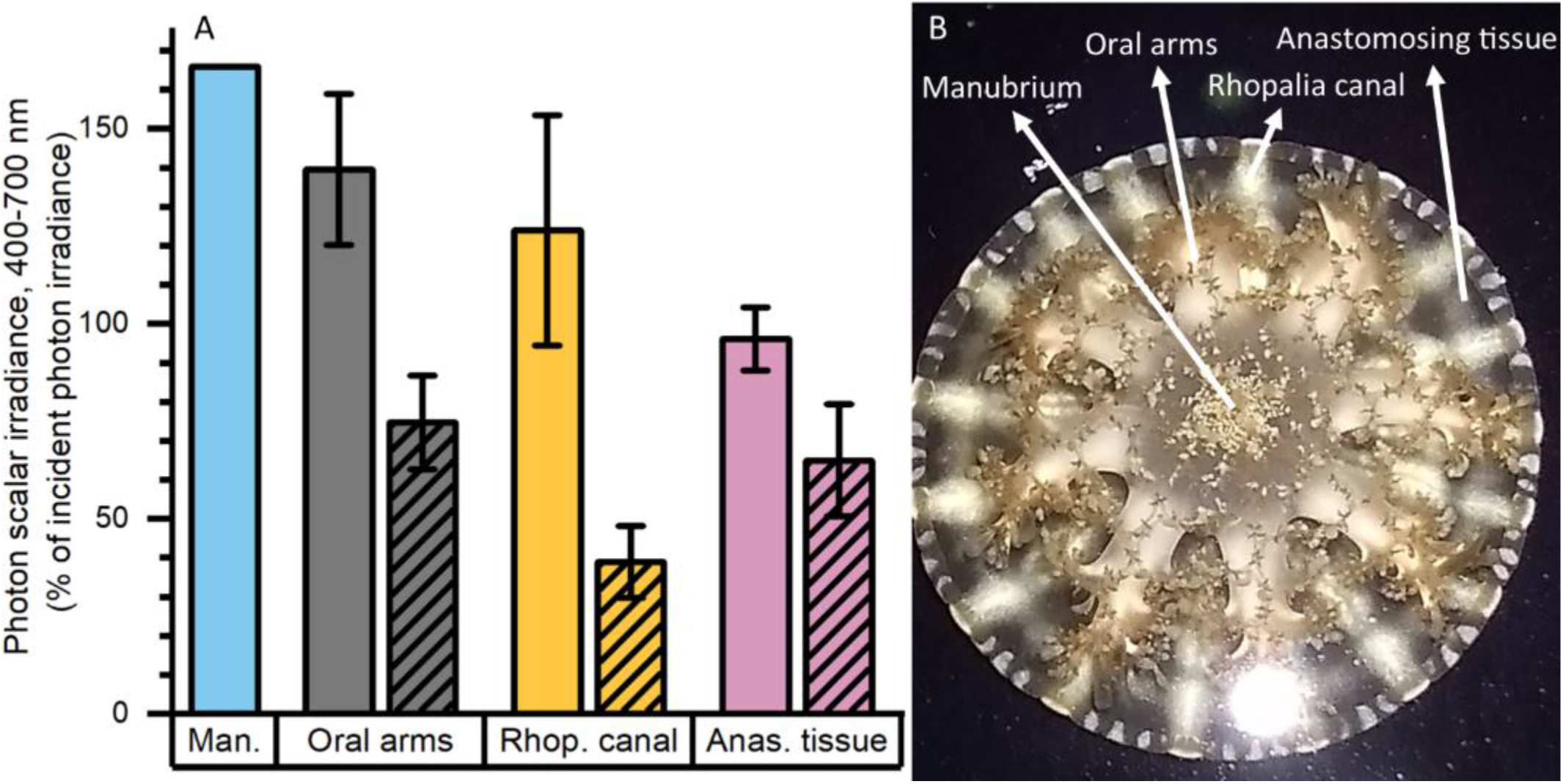
Normalized photon scalar irradiance (PAR, 400-700 nm) measured at (A) the manubrium center (Man.), oral arms, rhopalia canals (Rhop. canal) and anastomosing tissue (Anas. tissue) on a single adult medusa of *Cassiopea*. Bright colors indicate values measured above, and shaded colors the values below the medusa. (B) Overview of the specimen with exemplary positions for measurements (white arrows). Light intensities were mapped at the tissue-water interface (above and below the medusa) at respective anatomical structures (n = 7-17) using a submersible spherical quantum sensor (see methods), and normalized against the incident photon irradiance intensity measured at the same sensor positions without the medusae above a black surface. Note, only one measurement was performed above the manubrium center.

#### Spectral scalar irradiance & reflectance

Both spectral scalar irradiance and reflectance measurements required the medusa to be fixed in place on a large rubber stopper using hypodermic needles, and placed in a container filled with seawater. Incident light was provided vertically from above with a fiber-optic halogen lamp (KL-2500 LCD, Schott GmbH, Germany) fitted with a collimating lens. Spectral scalar irradiance was measured using a fiber-optic scalar irradiance microprobe (spherical tip diameter ~100 µm; Rickelt et al., 2016). Spectral reflectance was measured using a fiber-optic field radiance probe (400 μm diameter; Kühl, 2005). Both sensors were connected to a fiber-optic spectrometer (USB2000+; Ocean Optics, USA), and spectral information was acquired using SpectraSuite software (Ocean Optics, USA). Sensors were mounted on a manual micromanipulator (MM33, Märzhäuser Wetzlar GmbH, Germany) attached to a heavy-duty stand to facilitate precise positioning of sensors on the medusa. Sensors were positioned visually using a dissection microscope, while ensuring oral arms and other anatomical features did not shade the tissue studied. The spectral reflectance probe was positioned by first moving the sensor to the surface of the medusa, then moving the sensor 1 mm away. Similarly, the scalar irradiance probe was first positioned to touch the surface tissue of the medusa (depth = 0 µm) and was then moved into the medusa in vertical steps of 200 µm. Both sensors were held by the micromanipulator in an angle of 45° relative to the incident light. The recorded spectral scalar irradiance was normalized against the incident, downwelling spectral irradiance measured over the black cloth covering the inside of the container relative to each measuring position on the medusae. The recorded spectral reflection spectra were normalized against a reference spectrum measured with an identical setup over a 99% white reflectance standard (Spectralon, Labsphere) in air.

### Microscale measurements of O_2_ and pH

#### General setup

For all O_2_ and pH measurements, individual medusae were placed in a cylindrical plexiglass chamber with aerated, filtered ASW. Medusae were illuminated with a fiber-optic halogen lamp (KL-1500 LED, Schott GmbH, Germany) equipped with a collimating lens, and providing defined levels of incident photon irradiance (0, 42, 105, 160, 300, and 580 µmol photons m^−2^ s^−1^; 400-700 nm), as measured for specific lamp settings with a calibrated photon irradiance meter equipped with a spherical micro quantum sensor (ULM-500 and US-SQS/L, Heinz Walz GmbH, Germany).

Concentration profiles and dynamics were measured above and inside anastomosing bell tissue of 3 small (25-48 mm) and 2 large (63-67 mm) medusae. The microsensor tip was carefully positioned at the tissue surface-water interface while watching the tissue surface under a dissection microscope. For profiling, the sensor was inserted in vertical steps of 100 µm until reaching the approximate center of the mesoglea, relative to the subumbrella and exumbrella bell epidermis.

#### Oxygen measurements

The O_2_ concentration measurements were done using a robust fiber-optic O_2_ optode (OXR230 O_2_ sensor, 230 µm diameter tip, < 2 s response time; PyroScience GmbH, Germany) connected to a fiber-optic O_2_ meter (FireSting, PyroScience GmbH, Germany). The sensor was linearly calibrated in O_2_-free and 100 % air saturated seawater at experimental temperature and salinity. The sensor was mounted slightly angled relative to the incident light (to avoid shadowing) on a motorized micromanipulator (MU1, PyroScience GmbH, Germany) attached to a heavy-duty stand, facilitating visual sensor positioning with a dissection microscope. Data acquisition was done using the manufacturer’s software (Profix, PyroScience GmbH, Germany). Measurements were performed at room temperature (22°C) in a dark room under defined light conditions, and the temperature was continuously recorded in the experimental chamber with a submersible temperature sensor (TSUB21, PyroScience GmbH, Germany) connected to the O_2_ meter.

For light-dark measurements, the O_2_ sensor tip was inserted approximately midway between sub- and exumbrella epidermis (depth ranging from ~800 to 3000 µm from the surface depending on the specimen size). Once the depth of interest was reached, the local O_2_ concentration dynamics were measured at 10 s intervals during experimental manipulation of the light. Local rates of net photosynthesis and post-illumination respiration were estimated from the measured linear slope of O_2_ concentration versus time measurements (using linear fits in the software OriginPro 2020b) under light and dark conditions, respectively. The local gross photosynthesis was then estimated as the sum of the absolute values of net photosynthesis and respiration.

The O_2_ concentration depth profiles were recorded at vertical depth intervals of 100-200 µm during constant light or darkness. The measurements were recorded once the O_2_ concentration reached a steady-state at each depth, starting from deepest inside the tissue and moving the sensor up until the sensor tip was retracted up into the overlaying turbulent water column (100 % air saturation).

#### pH measurements

pH was measured using a pH glass microelectrode (PH-100, 100 µm tip diameter, response time < 10 s, Unisense, Denmark) combined with an external reference electrode and connected to a high impedance mV-meter (Unisense, Denmark). A motorized micromanipulator (MU1, PyroScience GmbH, Germany) was used for positioning of the sensor, and data were recorded using SensorTrace Logger software (Unisense, Denmark). Prior to measurements the sensor was calibrated at room temperature using IUPAC standard buffer solutions at pH 4, 7, and 9 (Radiometer Analytical, France). Dynamic changes in mesoglea pH levels were measured during experimental dark-light transitions and *vice versa*. pH depth profiles were measured under constant light or darkness, and were recorded from the middle of the bell and up into the water column in steps of 200 µm, until the sensor tip was retracted into the overlying turbulent water column (pH level of 8.1-8.4).

## Results

### Light microenvironment

#### Photon scalar irradiance

Macroscopic mapping of integral photon scalar irradiance of photosynthetically active radiation (PAR, 400-700 nm) in 4 distinct regions of an intact, living *Cassiopea* sp. revealed a heterogeneous light field across the medusa tissue facing the incident light as well as across the tissue. The manubrium center experienced the highest scalar irradiance reaching 166 % of incident photon irradiance (Figure 1). Oral arms experienced a photon scalar irradiance reaching 140 ± 19 % of incident photon irradiance, while light availability just below the oral arms was significantly lower (75 ± 12 % of incident photon irradiance). The light levels of rhopalia canals reached 124 ± 30 % of incident photon irradiance and light levels of anastomosing tissue reached 96 ± 8 % of incident photon irradiance on the subumbrella side. On the exumbrella side, more light penetrated the anastomosing tissue (65 ± 14 % of incident photon irradiance), while the least amount of light penetrated through the rhopalia canals (39 ± 9 % of incident photon irradiance).

#### Reflectance

The oral arms and rhopalia canals of *Cassiopea* reflected more light than the manubrium and anastomosing tissue, with oral arms reflecting 9.5 %, rhopalia canals 7 %, the manubrium 4.8 %, and anastomosing tissue 3 % of incident irradiance (Figure 2).

**Figure 2.**
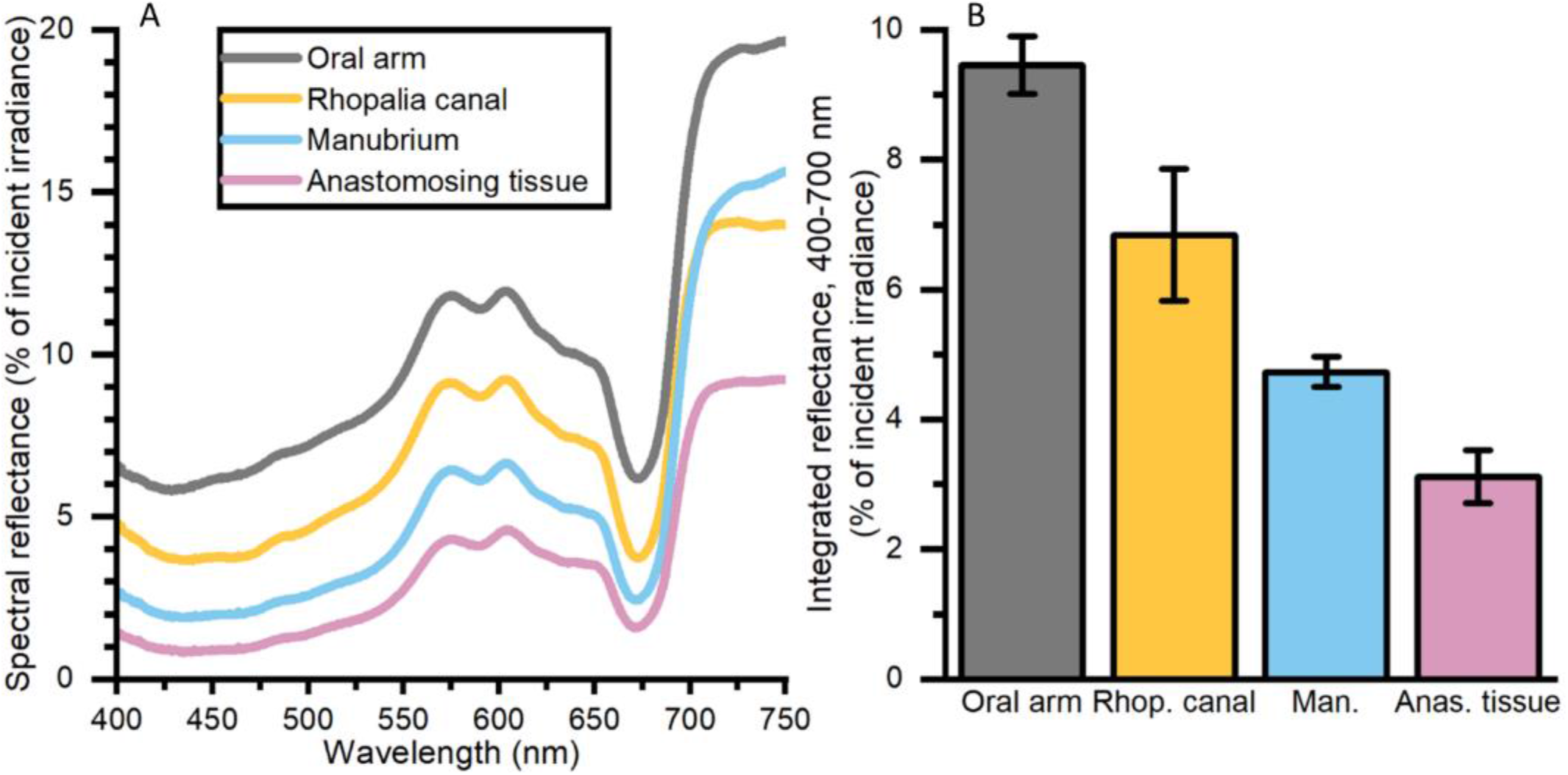
Spectral reflectance (in % of incident irradiance) (A) measurements, and integrated reflectance (400-700 nm) (B) averaged for 4 AOIs depicting differing morphological tissue regions of one large *Cassiopea* medusa. Data represent mean ±SEM (n = 3 for each tissue type). For increased clarity standard errors are not shown in panel A, but are reflected in the ±SEM shown for integrated values in panel B.

#### Spectral scalar irradiance

Depth profiles of spectral scalar irradiance in live *Cassiopea* tissue showed spectral absorption signatures of chlorophyll (Chl) *a* (430–440, 675 nm), Chl *c* (460, 580-590, 635 nm), and peridinin (480–490 nm), throughout the anastomosing tissue and near rhopalia canals (Figure 3). Spectral scalar irradiance at the tissue-water interface (0 µm) revealed a local enhancement near the rhopalia canals, while light was more readily absorbed on anastomosing tissue. Depth profiles show a steady attenuation of light within the first 400 µm of the bell near rhopalia canals (Figure 3A). In contrast, scalar irradiance was slightly enhanced at 200 and 400 µm in the anastomosing tissue, until a strong attenuation was observed at 600 µm depth (Figure 3B).

**Figure 3.**
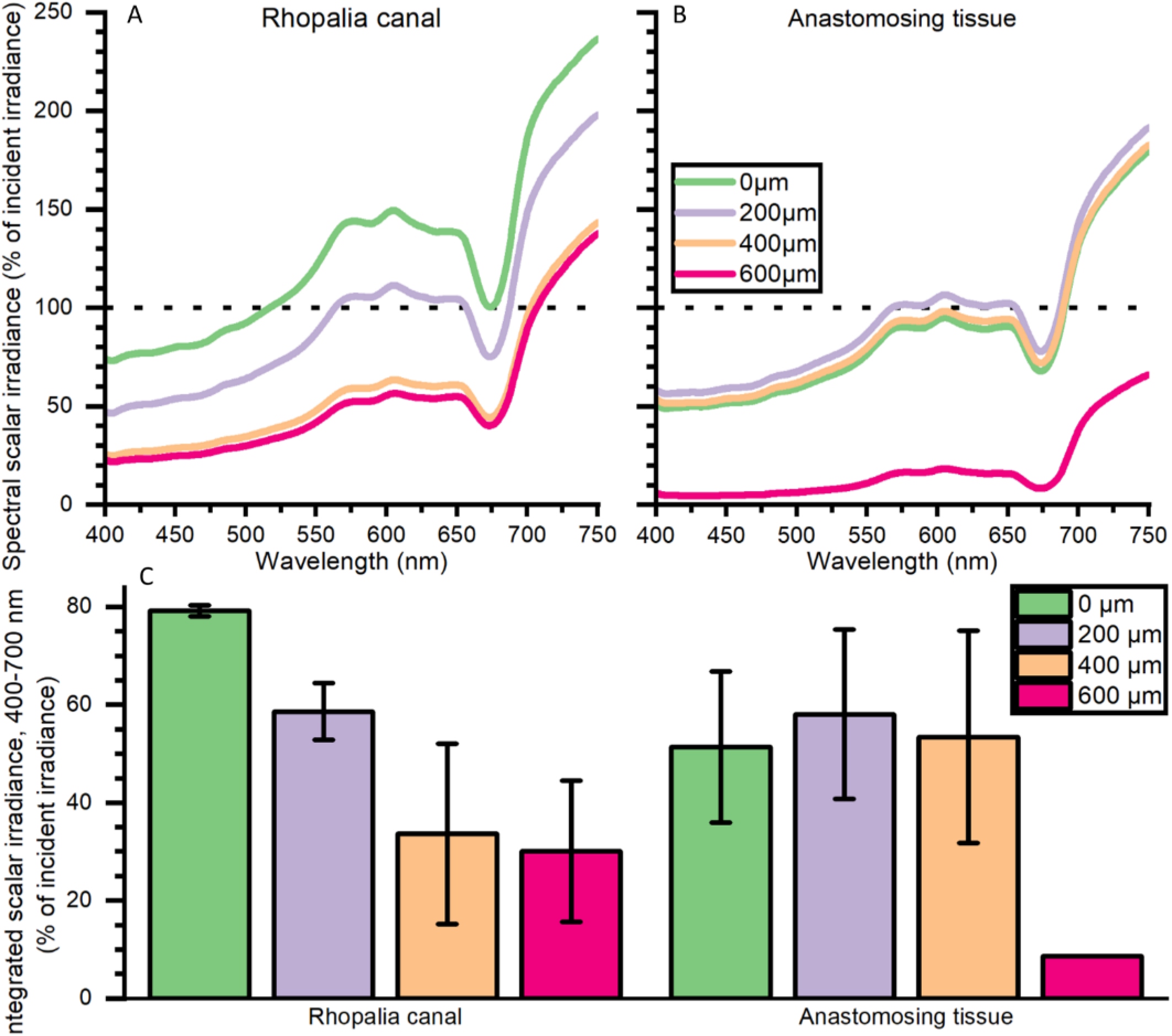
Spectral scalar irradiance (in % of incident irradiance) measured near the subumbrella rhopalia canal (A) and anastomosing tissue (B), and integrated scalar irradiance (400-700 nm) (C). Data represent mean ±SEM of 2 rhopalia canals and 3 anastomosing tissue areas (except for the 600 μm). For increased clarity standard errors are not shown in panel A and B, but are reflected in the ±SEM shown for integrated values in panel C.

### Chemical microenvironment

#### Oxygen depth profiles

The O_2_ concentration depth profiles were measured in a *Cassiopea* sp. medusa (45 mm Ø) under a saturating photon irradiance of 300 µmol photons m^−2^ s^−1^ (Figure 4). The O_2_ concentration increased strongly in deeper parts of the bell tissue below 1 mm depth, rising from 315 (± 21) to 445 (± 40) µmol O_2_ l^−1^ at a depth of 3.7 mm. A maximal O_2_ concentration was measured at 4 mm, reaching ~500 µmol O_2_ l^−1^ (not shown in figure).

**Figure 4.**
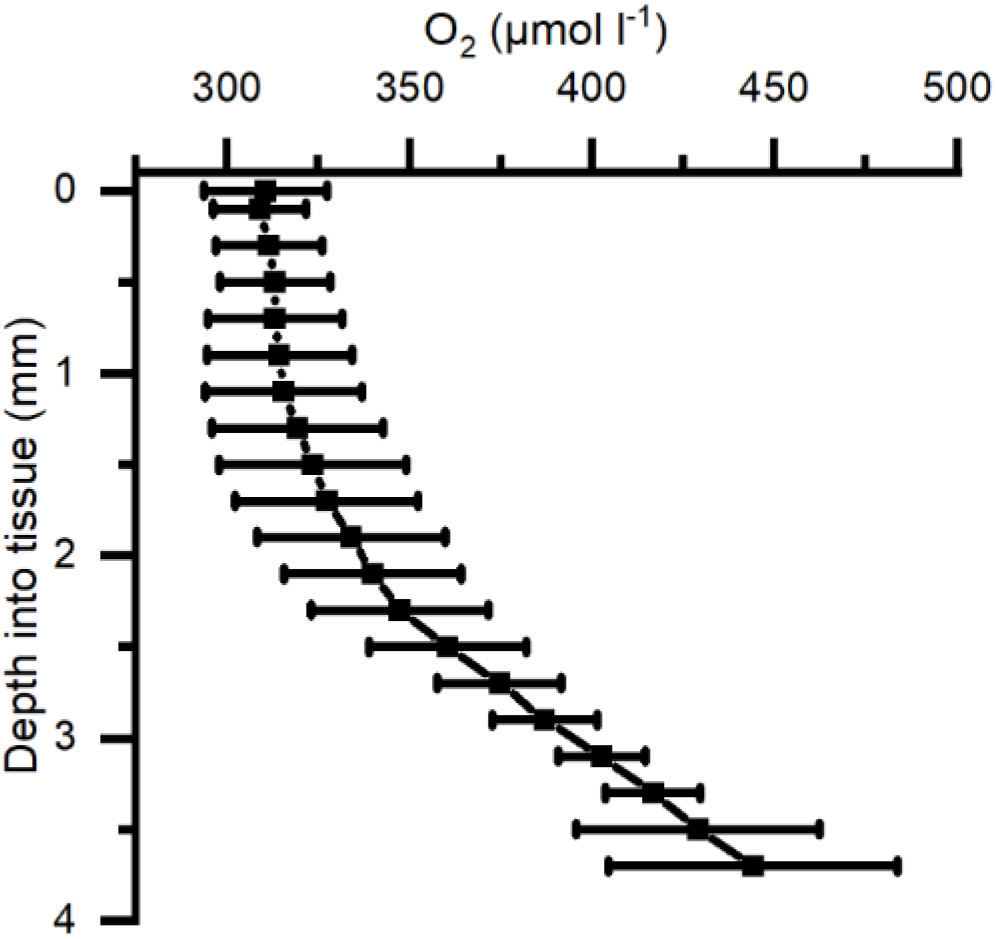
Average of depth profiles of O_2_ concentration in *Cassiopea* tissue measured under a photon irradiance (400-700 nm) of 300 μmol photons m^−2^ s^−1^ in the bell of a medusae. n = 3 biological replicates with 1 profile each.

Similarly, O_2_ depth profiles were measured in darkness in 3 different medusae (38-67 mm Ø), with either 0, 15, or 50 min of darkness prior to the profile (Figure 5). Profiles reveal high O_2_ depletion in the top 1 mm layer of the bell with increasing length of dark incubation, while measurements deeper into the bell showed a less dramatic decrease of O_2_ concentration in the mesoglea of the medusae.

**Figure 5.**
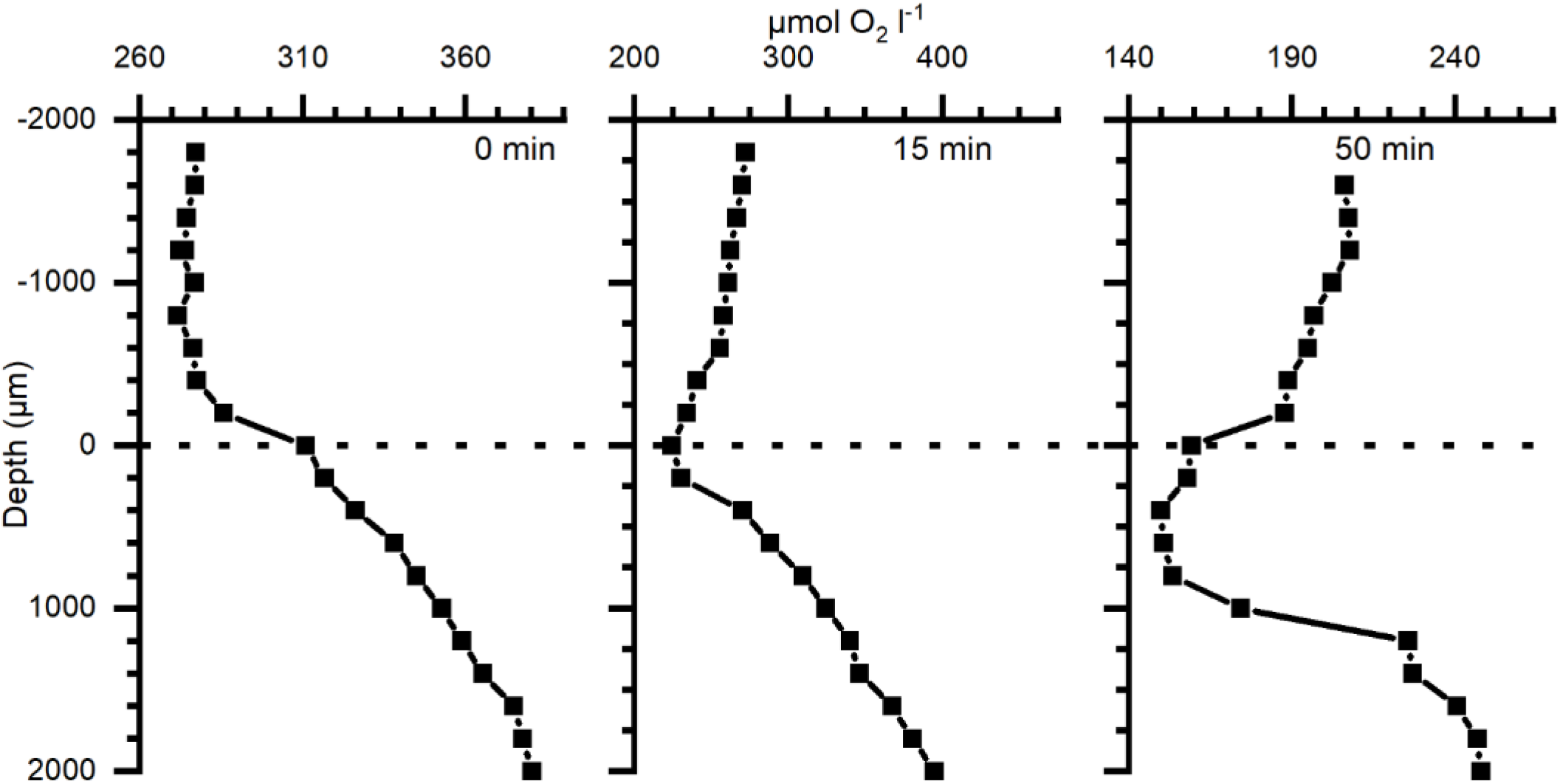
Depth profiles of O_2_ concentration in bell tissue of 3 different *Cassiopea* specimens (38-67 mm Ø) incubated in darkness for 0, 15, and 50 min, respectively (38, 67, and 63 mm Ø individual, respectively), prior to measurements. Zero depth indicates the tissue-water interface. Note differences in the x-axis scales.

#### Light-dependent oxygen dynamics

The O_2_ dynamics measured approximately in the middle of the mesoglea between the sub- and exumbrella epidermis showed a delayed response to changes in light in large individuals, while the response in smaller medusae appeared immediately after a light-dark switch or *vice versa* (Figure 6). The O_2_ concentration measured in 2 large and 3 small medusae showed that the O_2_ content continued to increase in the mesoglea of large medusae several minutes after light was turned off, indicative of diffusive supply from the surrounding tissue with a higher symbiont density and/or photosynthetic activity prior to onset of darkness (Figure 7).

**Figure 6.**
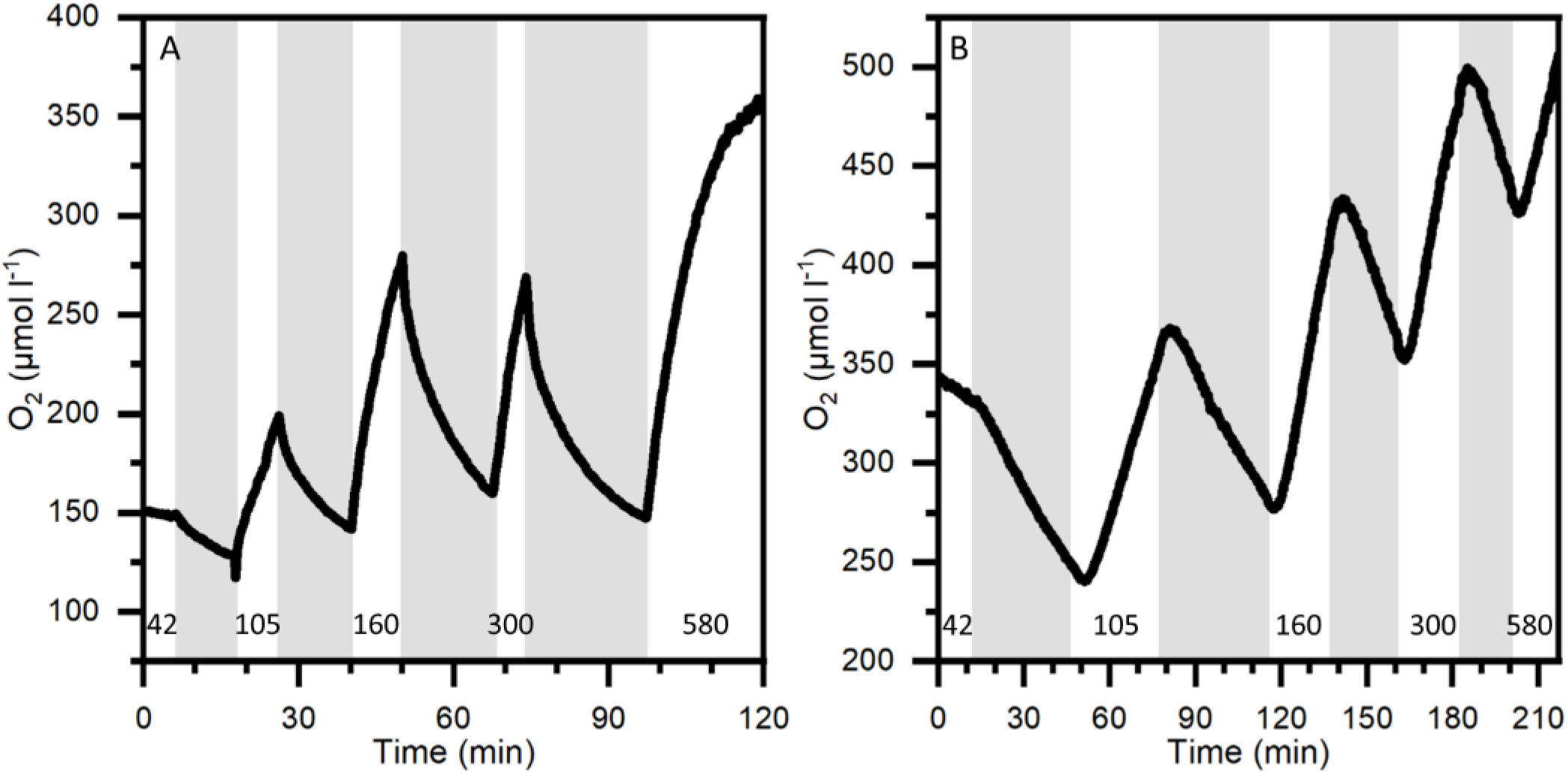
O_2_ dynamics at 1 mm depth into the bell of a small medusa (A) and 3 mm into the bell of a large medusa (B) during light-dark shifts. Gray areas indicate dark shifts and white areas indicate photo periods of increasing incident photon irradiance (400-700 nm) noted with numbers in units of μmol photons m^−2^ s^−1^. Note differences in x- and y-axes.

**Figure 7.**
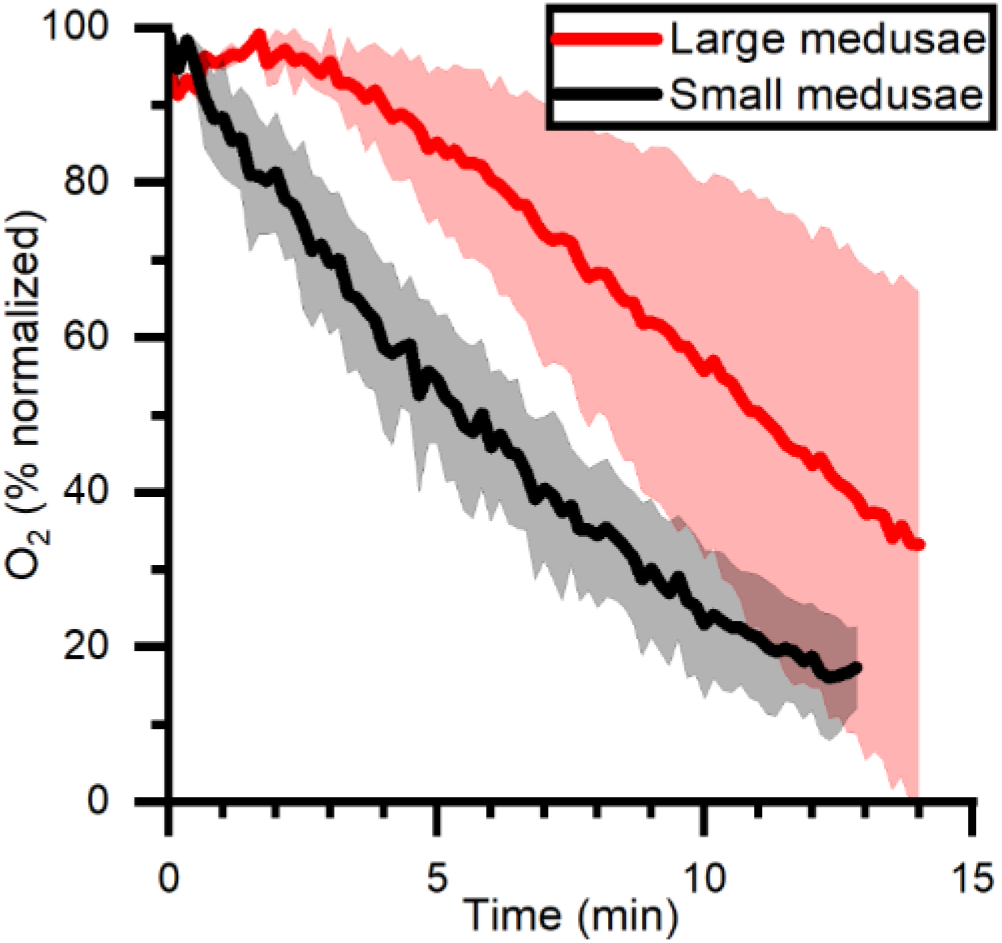
O_2_ dynamics measured in mid-bell mesoglea (roughly halfway deep in the bells) of large and small medusae from onset of darkness at 0 min, after a previous illumination with an incident photon irradiance (400-700 nm) of 580 μmol photons m^−2^ s^−1^. Data is normalized against maximum oxygen concentration measured in each medusa, and then averaged for large (n=2) and small (n=3) individuals ±SEM.

Locally measured photosynthesis versus photon irradiance curves showed that photosynthesis increased linearly with irradiance until saturation was approached at ~300 µmol photons m^−2^ s^−1^ (Figure 8). At saturating irradiance, net photosynthesis rates in bell tissue ranged from 0.08 µmol O_2_ l^−1^ s^−1^ in large individuals, to 0.18 µmol O_2_ l^−1^ s^−1^ in small medusae. Similarly, the estimated gross photosynthesis was roughly 2-fold higher in small medusae, at 0.26 μmol O_2_ l^−1^ s^−1^ *vs*. 0.14 μmol O_2_ l^−1^ s^−1^ in large medusae. Post-illumination respiration rates were 0.04-0.07 μmol O_2_ l^−1^ s^−1^ in large medusae regardless of illumination, while post-illumination respiration in small medusae increased to a maximum of 0.18 μmol O_2_ l^−1^ s^−1^ at maximum photon irradiance (580 μmol photons m^−2^ s^−1^). The respiration at saturating photon irradiance (300 μmol photons m^−2^ s^−1^) was 0.08 μmol O_2_ l^−1^ s^−1^ in tissue of small medusae.

**Figure 8.**
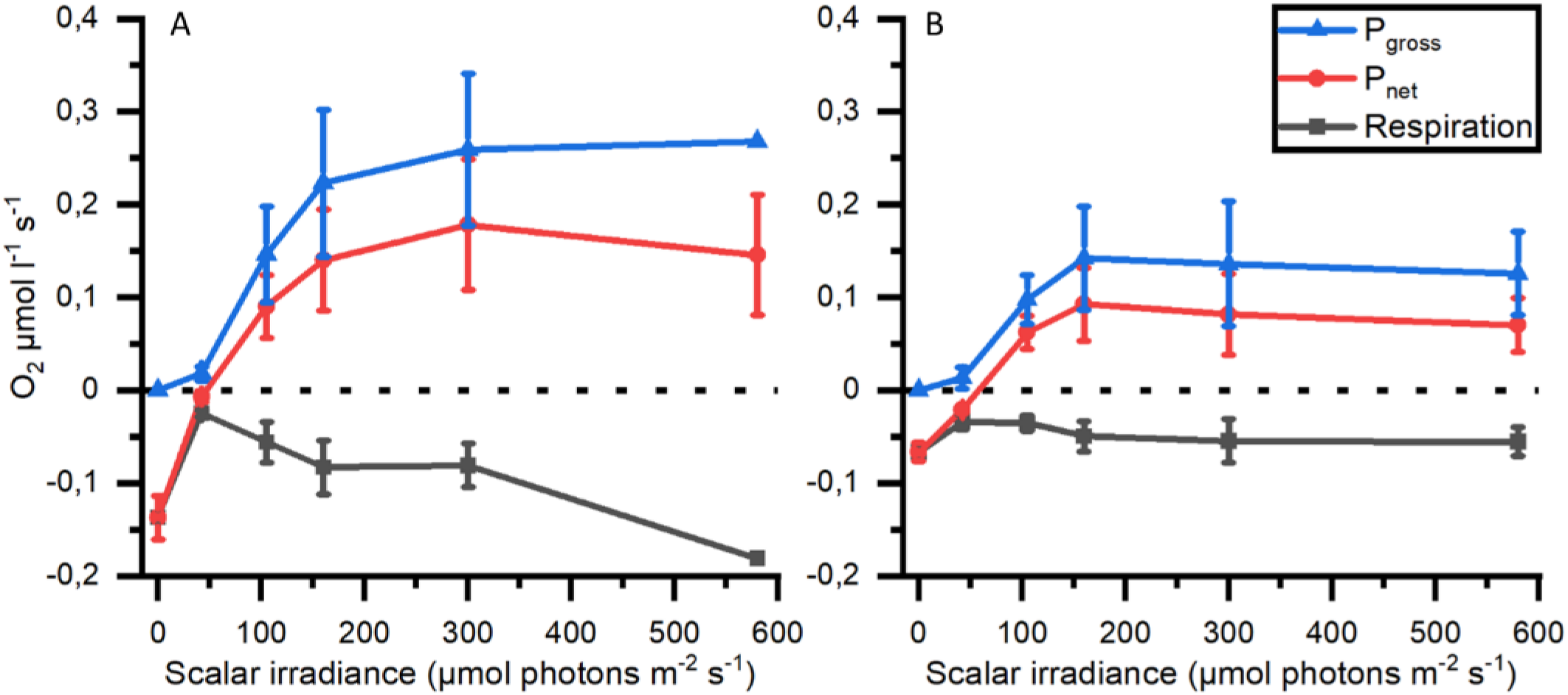
Respiration, net (P_net_) and gross photosynthesis (P_gross_) measured mesoglea of small (A; n=3) and large (B; n=2) medusae as a function of photon scalar irradiance (400-700 nm).

#### Spatio-temporal dynamics of pH

pH depth profiles measured after a combined 15 minutes light, 15 minutes darkness incubation showed that pH dropped rapidly to below ambient water pH at the bell tissue-water interface (0 μm) in both the large and small medusae (Figure 9). pH then increased rapidly again within the first 1000 μm of the bell tissue (to above ambient water pH), and kept increasing to a maximum depth of 4000 μm to a final pH of ~8.8 for both specimens. The pH dynamics over experimental light-dark shifts showed a similar pattern to what was observed with O_2_ dynamics: pH changed relatively quickly in the small medusa, while a much longer delay (usually > 10 min after the light change) was observed with the large medusa (Figure 10).

**Figure 9.**
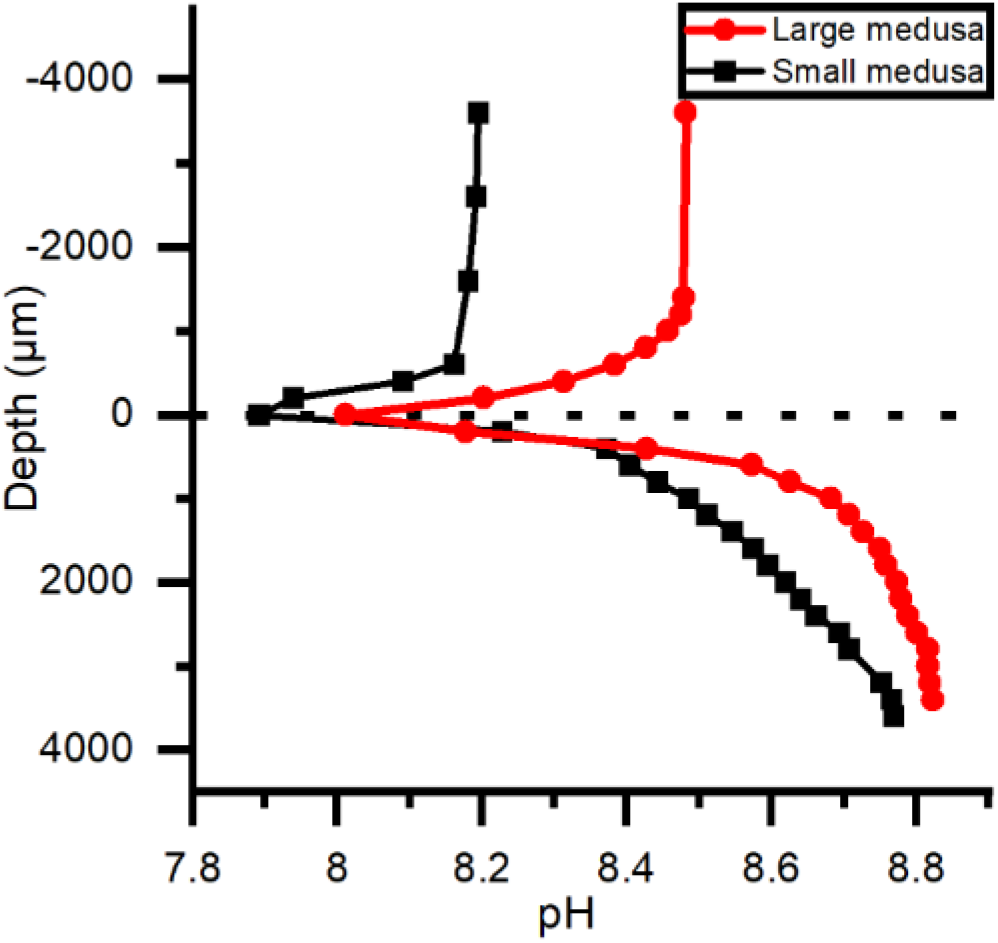
pH depth profiles measured in the bells of a small and large medusae. Both individuals were kept under an incident photon irradiance (400-700 nm) of 580 μmol photons m^−2^ s^−1^ light for >15 minutes and then in darkness for ~15 minutes before profiles were done.

**Figure 10.**
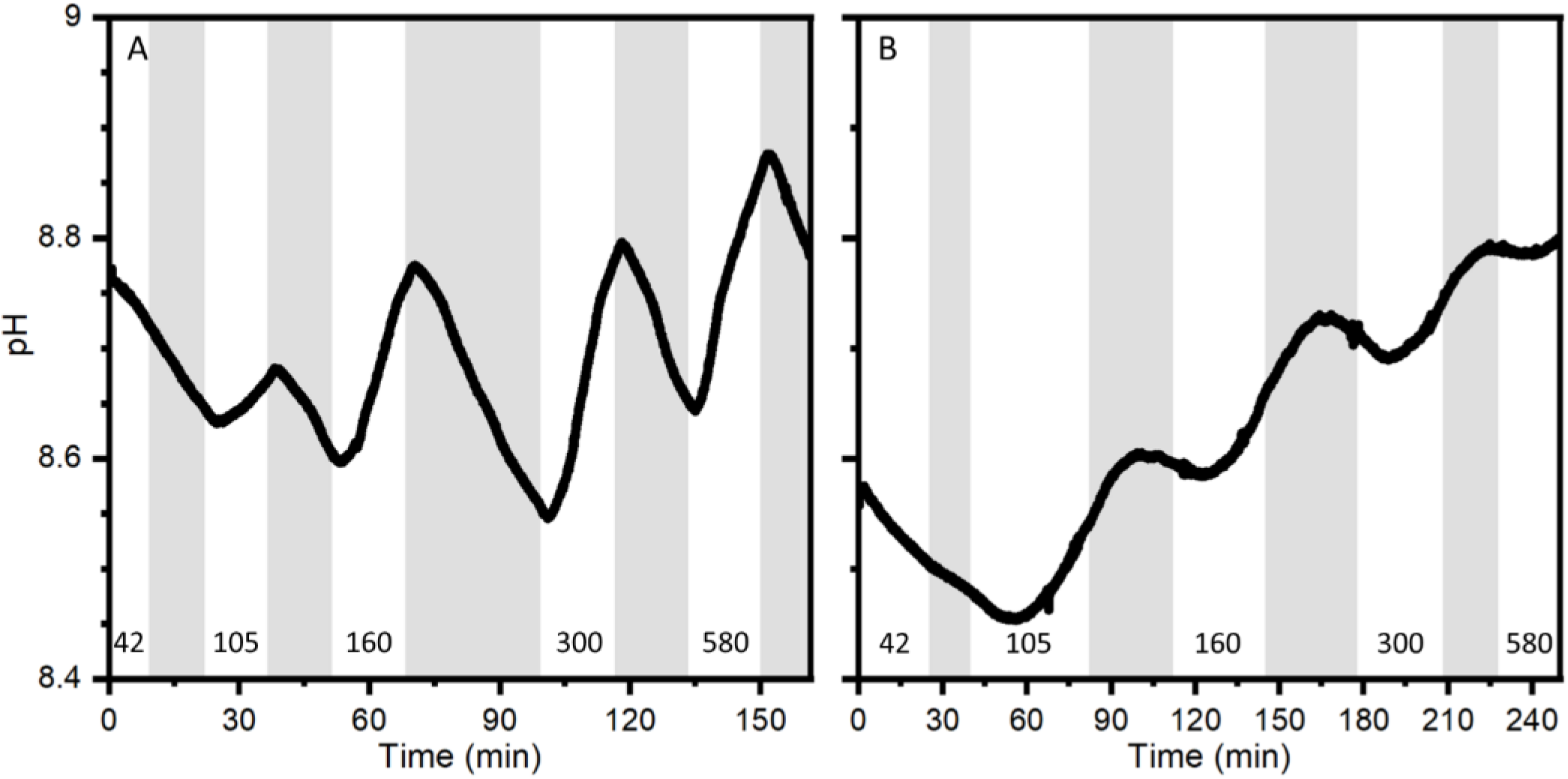
pH dynamics measured with a pH microsensor positioned at a depth of 1.6 mm into the bell of a small medusa (A) and 2 mm into the bell of a large medusa (B) during experimental light-dark shifts. Gray areas indicate dark shifts and white areas indicate photo periods of increasing incident photon irradiance (400-700 nm) noted with numbers in units of μmol photons m^−2^ s^−1^. Note difference in x-axes.

## Discussion

### *Cassiopea* harbors optical microniches

Light availability in the host is a key factor for an efficient photosymbiosis. It hast been shown that host morphology and tissue plasticity, host pigmentation, and symbiont density can modulate the scattering and absorption of photons in tissue of reef building corals, creating optical microniches in a colony (D’Angelo et al., 2008; Wangpraseurt et al., 2014b, 2019; Bollati et al., 2021); and we note that the coral skeleton is also important for the coral light microenvironment (e.g. Enriquez et al., 2005; Enríquez et al., 2017). In *Cassiopea*, we found distinct spatial heterogeneity over the surface tissue in various regions of an adult medusa (Figure 1). Local photon scalar irradiance was highest near apical parts of the animal, while it dropped by 30-80 % underneath the same anatomical structures (Figure 1A). The largest decrease was observed between measurements on the subumbrella and exumbrella epidermis along the rhopalia canal, which dropped by 80 % on the exumbrella side. Additionally, spatial heterogeneity was observed over the subumbrella bell tissue, with rhopalia canals experiencing on average ~30 % higher photon flux compared to the anastomosing tissue right next to it.

Depth profiles of spectral scalar irradiance measured in tissue close to rhopalia canals and in anastomosing tissue showed that light scattered differently within the upper 600 μm of the two regions (Figure 3). Profiles in anastomosing tissue showed small variations in spectral scalar irradiance measured in the top 400 μm of this region (Figure 3B). Such small variations indicate the presence of a dense symbiont population spread out in the top 400 μm of the mesoglea, between the subumbrella epidermis and the gastric network (Estes et al., 2003). Similar studies on corals have shown that light can be scattered by host tissue and symbionts in the surface layers, enhancing the chance of photon absorption (Wangpraseurt et al., 2012, 2014a; Jacques et al., 2019). In contrast, we observed a strong light attenuation over the first 400 μm of the tissue near rhopalia canals, with an initial enhancement of local scalar irradiance that quickly attenuated to levels below what was observed at the same depth in anastomosing tissue (Figure 3B). *Cassiopea* contain distinct white striated patterns in dense layers along the rhopalia canals in the bell and under oral arms (see white patches in Figure 1B) (Bigelow, 1900). While little is known about host pigments in *Cassiopea* and other rhizostome jellyfish (Hamaguchi et al., 2021; Lawley et al., 2021), our reflection data (compare e.g. rhopalia canal with anastomosing tissue in Figure 2) suggests that the white pigmented tissue can play a role in scattering and reflection of light in *Cassiopea* medusae, similar to host pigments and the skeleton in reef building corals (Wangpraseurt et al., 2014a; Jacques et al., 2019). More detailed analyses are required to understand the full potential of light-modulating host pigments in *Cassiopea*, and how the holobiont might respond and grow under various light regimes.

### *Cassiopea* harbors its photosymbionts in a buffered chemical microenvironment

We investigated the light-driven dynamics of O_2_ and pH on the surface and inside the bell tissue of *Cassiopea* medusae. Depth profiles of O_2_ concentration measured at saturating irradiance showed the presence of a diffusive boundary layer (DBL) over the bell tissue (Figure 5). In light, the O_2_ concentration increased towards the bell tissue-water interface and continued to increase as the sensor penetrated deeper into the tissue and mesoglea (Figure 4). We measured to a maximum depth of 4 mm into the mesoglea reaching an O_2_ concentration of roughly 2-fold that of the surrounding seawater concentration. A similar study, measuring O_2_ in cut-off oral arms of *Cassiopea* sp., found that O_2_ concentration was highest near symbiont populations (i.e. near the epidermis), while it decreased to a more constant level deeper into the oral arm mesoglea (Arossa et al., 2021). However, our measured depth profiles in bell tissue did not show any indication that O_2_ buildup would stagnate or decrease at this point, suggesting even higher concentrations might be possible in the bell mesoglea relative to oral arms. Passive accumulation of O_2_ has been reported in non-symbiotic jellyfish like *Aurelia labiata* (Thuesen et al., 2005), which was found to build up O_2_ in the mesoglea when in O_2_-rich water, indicating that the mesoglea can act as a natural reservoir for O_2_ in jellyfish. However, similar to measurements in oral arms of *Cassiopea* sp. (Arossa et al., 2021), measured depth profiles of O_2_ concentrations in non-symbiotic jellyfish like *Aurelia* appear to have the highest O_2_ concentration near the tissue-water interface, with a steady decline towards the center of the bell (Thuesen et al., 2005). In contrast our measurements indicate that the presence of photosynthetic endosymbionts harbored within amoebocytes in the mesoglea (Colley and Trench, 1985; Medina et al., 2021) leads to internal O_2_ production and accumulation in deeper tissue layers during photo periods (Kühl et al., 1995; Arossa et al., 2021).

The diffusion of O_2_ through cnidarian tissue and mesoglea has received little attention, however, Brafield and Chapman (1983) determined an O_2_ diffusion coefficient of 7.69×10^−6^ cm^2^ s^−1^ (Fick’s law) in the mesoglea of the sea anemone *Calliactis* sp. Such a low diffusion coefficient in the mesoglea (as compared to seawater) will impede mass transfer, especially in cnidarians with a particularly thick mesoglea like jellyfish and sea anemones, and is probably a key factor in the observed buffering of O_2_ dynamics in the *Cassiopea* sp. bell. We further investigated the dynamics of O_2_ in darkness and found that the top 1 mm layer of *Cassiopea* bell tissue turned from a net O_2_ source into a sink (Figure 5). Measurements in the DBL showed a switch from a net export from the tissue into the surrounding seawater to a diffusive import of O_2_ from the seawater within the first 15 min (compare DBLs in Figure 5). A more pronounced depletion of O_2_ was observed in the top 1 mm of the bell after 50 min darkness. The higher O_2_ consumption near the subumbrella epidermis probably reflects the presence of abundant musculature required for bell pulsation and motility of jellyfish (Blanquet and Riordan, 1981; Thuesen et al., 2005; Aljbour et al., 2017), as well as the presence of a dense population of endosymbionts (Estes et al., 2003; Lampert, 2016). Both have previously been ascribed to heavy diel fluctuations of O_2_ measured in *Cassiopea* oral arms (Arossa et al., 2021). However, the O_2_ concentration remained high at depths deeper than 1 mm into the bell even after 50 min of darkness, reflecting a lower cell density and diffusive transport of O_2_.

In the present study, experimental light-dark shifts performed on intact small (< 5 cm) and large (> 6 cm) *Cassiopea* sp. showed that the O_2_ concentration in small medusae with a (relatively) thin bell of 2 mm thickness changed almost immediately in response to light-dark shifts similar to observed O_2_ dynamics in corals and dissected oral arms of *Cassiopea* (Figure 6A) (Kühl et al., 1995; Arossa et al., 2021). Unlike the fast response observed in small medusae, larger medusae with a thicker bell tissue showed a much slower response of O_2_ levels to changes in light (Figure 6B). In fact, detailed O_2_ dynamics measured in several large and small medusae revealed a consistent pattern where O_2_ would continue to build up in the thick mesoglea of larger medusae for a few minutes after onset of darkness (Figure 7). These observations indicate that O_2_ generated in other regions with higher photosynthetic activity due to higher light levels and/or higher symbiont density can diffuse into and accumulate in the mesoglea, from where it is not efficiently exchanged with the surrounding water due to the relatively low O_2_ diffusivity in mesoglea (Brafield and Chapman, 1983).

Furthermore, an inverse relationship between medusa size and photosynthetic rate have previously been reported (Verde and McCloskey, 1998). We observed a similar inverse relationship, with smaller medusae on average reaching roughly 2-fold higher photosynthetic rates (net and gross) at photon irradiances above 200 μmol photons m^−2^ s^−1^ (Figure 8). Post-illumination respiration also increased in small individuals, while larger medusae did not show changes to respiration after illumination. The combined effect of the diffusive properties of mesoglea, the relative thickness of the mesoglea, and the overall size of the animal might together explain why the O_2_ dynamics is more pronounced in medusae (and other cnidarians) with a thin mesoglea like in juvenile *Cassiopea* bell or oral arms, while the O_2_ dynamics in medusae with a thick mesoglea is much more buffered.

Consistent with the measured O_2_ dynamics, we found that pH changes in the mesoglea were affected by changes in light, because pH increases due to photosynthetic carbon fixation in the light and decreases during darkness due to respiration (Figure 10). This relationship between photosynthesis, respiration, and pH seems prevalent in symbiotic cnidaria (e.g. Kühl et al., 1995; de Beer et al., 2000; Chan et al., 2016; Klein et al., 2017; Arossa et al., 2021), largely driven by the shifting equilibrium between carbonate species (i.e.; CO_2_, H_2_CO_3_, HCO_3_^−^, CO_3_^2−^) and the balance between photosynthesis and respiration. de Beer et al. (2000) found that pH in coral tissue followed the trend of O_2_ during experimental light-dark shifts but with a delayed response (seconds to minutes), and attributed the delay to hypothetic processes, such as proton pumps and other similar cross-tissue transport, that would buffer pH in coral polyps (Palmer and Van Eldik, 1983). External buffering has also been reported in polyps of *Cassiopea*, where Klein et al. (2017) found that polyps retained their internal pH at ambient water pH during the night, and only symbiotic polyps would increase their internal pH during photoperiods due to a shifting carbonate equilibrium. We observed a similar, but much longer (>10 min) delay before a pH change was observed in the mesoglea of large medusa after light-dark shifts (Figure 10B). While external buffering is likely to occur in medusae of *Cassiopea* as well, the pH dynamics inside the mesoglea is probably more affected by diffusive transport phenomena. Depth profiles done in both small and large medusae thus show that within 15 min of darkness the top 1 mm of the bell tissue became more acidic relative to ambient water pH (0.3-0.5 pH lower; Figure 9), while pH in deeper tissue layers (>1000 μm into the bell) remained alkaline and above ambient water pH. This strongly suggest that the mesoglea has a buffering effect on pH in the bell tissue of *Cassiopea*.

The buffering capacity of the mesoglea could be of benefit for both host and symbionts. A reservoir of O_2_ can e.g. act as a steady supply of O_2_ to both host musculature, needed for bell pulsation, and to symbionts during dark periods or exposure to hypoxia (Thuesen et al., 2005). Similarly, buffering pH could lower the possibility of the holobiont experiencing cellular acidosis (Smith and Raven, 1979; Gibbin et al., 2014) that would otherwise disrupt cell-function (Madshus, 1988). Thus, symbionts harbored in amoebocytes in the *Cassiopea* mesoglea may exist in a more stable ecological niche as compared to algae in corals that are more prone to rapid chemical dynamics. We speculate that the buffering of the chemical microenvironment in the mesoglea of *Cassiopea*, might also be reflected in the fact that *Cassiopea* medusae generally seem to only engage with a specific type of Symbiodiniaceae. Indeed, specific strains of Symbiodiniaceae have been speculated to be favored by different hosts in symbiotic cnidarians (Schoenberg and Trench, 1980; Biquand et al., 2017), including *Cassiopea* (Colley and Trench, 1983; Fitt, 1985). While specificity is generally attributed to a combination of symbiont cell size and a hospitable host microenvironment (Biquand et al., 2017), Fitt (1985) found a correlation between symbiont cell size and respective photosynthesis and respiration rates, and proposed that only specific symbiont strains are able to establish symbiosis due to metabolic rates matching the hosts specific microenvironment. As such, the *Cassiopea*-Symbiodiniaceae symbiosis may be successful even in extreme environments as *Cassiopea* is capable of maintaining less stress-tolerant species due to the buffering nature of the *Cassiopea* chemical microenvironment.

## Summary

In comparison to corals, *Cassiopea* shares similar optical properties to that of reef-building coral tissue. Both macroscale host anatomy and microscale structures play a role in modulating the internal light field experienced by the symbionts via light scattering. Precisely how these structures affect symbiont photosynthesis remains to be explored further. Furthermore, our microsensor measurements indicated a buffering of chemical dynamics in the thick mesoglea matrix of *Cassiopea* sp. medusae, suggesting that the internal physico-chemical microenvironment of the holobiont remains more constant when experiencing abrupt changes in light conditions. This is in strong contrast to the rapid dynamics seen in coral tissue, which has a much thinner mesoglea and where the endosymbionts are found within endoderm cells. We hypothesize that the stabilization of the internal host microenvironment can be beneficial to the holobiont during unfavorable external environmental conditions such as hypoxia, where stored O_2_ in the mesoglea might act as an important reserve for keeping internal homeostasis. This may also be key to the apparent success of *Cassiopea* to invade and persist coastal tropical habitats with strong environmental fluctuations, that often preclude coral colonization. However, further studies are required to determine the effect of the buffering capacity of large individuals in combination with true environmental stressors over short and long periods.

## Author Contributions

MCM, NHL, ET, and MK designed the experiment. Data acquisition was done by MCM, ET, and MK. Data and statistical analyses were carried out by MCM, NHL, ET, and MK. All authors contributed to writing and editing of the manuscript.

## Acknowledgments

We thank Sofie Lindegaard Jakobsen for excellent technical assistance with keeping *Cassiopea* specimens used in this study.

